# Evidence for genetic correlations and bidirectional, causal effects between smoking and sleep behaviours

**DOI:** 10.1101/258384

**Authors:** Mark Gibson, Marcus R Munafò, Amy E Taylor, Jorien L. Treur

**Affiliations:** School of Experimental Psychology, University of Bristol, Bristol, UK; School of Experimental Psychology, University of Bristol, Bristol, UK; MRC Integrative Epidemiology Unit at the University of Bristol, Bristol, UK; UK Centre for Tobacco and Alcohol Studies, Bristol, UK; Population Health Sciences, Bristol Medical School, University of Bristol, Bristol, UK; NIHR Biomedical Research Centre at the University Hospitals Bristol NHS Foundation Trust and the University of Bristol, UK; School of Experimental Psychology, University of Bristol, Bristol, UK; MRC Integrative Epidemiology Unit at the University of Bristol, Bristol, UK

**Author notes:** Shared senior author (Amy E Taylor, Jorien L. Treur). Corresponding author: Jorien L Treur, School of Experimental Psychology, University of Bristol, 12a Priory Road, Bristol BS8 1TU, UK.

**Keywords:** smoking, sleep, chronotype, insomnia, genetic correlation, Mendelian randomization

## Abstract

**Introduction:** Cigarette smokers are at increased risk of poor sleep behaviours. However, it is largely unknown whether these associations are due to shared (genetic) risk factors and/or causal effects (which may be bi-directional).

**Methods:** We obtained summary-level data of genome-wide association studies of smoking (smoking initiation (*n*=74,035), cigarettes per day (*n*=38,181) and smoking cessation (*n*=41,278)) and sleep behaviours (sleep duration and chronotype, or ‘morningness’) (*n*=128,266) and insomnia (*n*=113,006)). Using LD score regression, we calculated genetic correlations between smoking and sleep behaviours. To investigate causal effects, we employed Mendelian randomization (MR), both with summary-level data and individual level data (n=333,581 UK Biobank participants). For MR with summary-level data, individual genetic variants were combined with inverse-variance weighted meta-analysis, weighted median regression and MR Egger regression methods.

**Results:** We found positive genetic correlations between insomnia and smoking initiation (*rg*=0.27, 95% CI 0.06 to 0.49) and insomnia and cigarettes per day (*rg*=0.15, 0.01 to 0.28), and negative genetic correlations between sleep duration and smoking initiation (*rg*=-0.14, -0.26 to -0.01) and chronotype and smoking cessation (*rg*=-0.18, -0.31 to -0.06). MR analyses provided strong evidence that smoking more cigarettes per day causally decreases the odds of being a morning person, and weak evidence that insomnia causally increases smoking heaviness and decreases smoking cessation odds.

**Conclusions:** Smoking and sleep behaviours show moderate genetic correlation. Heavier smoking seems to causally affect circadian rhythm and there is some indication that insomnia increases smoking heaviness and hampers cessation. Our findings point to sleep as a potentially interesting smoking treatment target.

## Introduction

Observationally, cigarette smoking is associated with poor sleep. Smokers take a longer time to fall asleep and are at higher risk of experiencing sleep disturbances^1^. While sleep duration is generally shorter in smokers, longer than average sleep duration is more common too^2,3^. One longitudinal study that followed substance naïve adolescents into early adulthood, found that erratic sleep patterns predicted smoking initiation^4^. In adults, transitioning from ‘adequate’ to ‘inadequate’ sleep duration over a period of five years predicted heavier smoking^5^ and pre-existing insomnia symptoms increased the likelihood of relapse after an attempt to quit smoking^6^. Chronotype – being a ‘morning’ versus an ‘evening’ person – has also been linked to smoking such that smokers are more likely to be an evening person^7^. This is in contrast to evidence suggesting that young adolescents with an evening chronotype show a lower odds of smoking initiation four to five years later^4^. The observational nature of the studies described here precludes strong conclusions about causality. Unravelling the nature of the relationship between smoking and poor sleep is important given the major health burden that both behaviours pose^8,9^.

Observational associations between smoking and sleep may reflect common risk factors. These could be environmental in nature, such as socioeconomic factors^10,11^, or genetic – twin and family studies have reported a moderate to high heritability for both smoking and sleep^2,12,13^. Genome-Wide Association (GWA) studies have identified genetic variants robustly associated with smoking initiation, number of cigarettes smoked per day and smoking cessation^14^, and more recently, sleep duration and chronotype^15^ and insomnia^16^. This knowledge of the genetic architecture of smoking and sleep allows us to investigate the degree to which genetic risk for both phenotypes overlaps. With Linkage Disequilibrium (LD) score regression, a genetic correlation between two phenotypes can be calculated such that ‘0’ reflects no overlap in genetic variants and ‘1’ reflects that all genetic variants perfectly overlap^17^. Hammerschlag and colleagues reported sizeable, positive, genetic correlations between smoking and insomnia complaints^16^, but whether smoking behaviour shows genetic overlap with sleep duration and chronotype is still unknown. Moreover, genetic correlations may also reflect causal relationships. If, for instance, smoking cigarettes causally increases insomnia, then genetic variants that underlie vulnerability for smoking will also be associated with insomnia.

Causal effects could operate in either direction, from smoking to poor sleep (possibly due to nicotine’s stimulating effects) and from poor sleep to smoking (such that cigarettes are used as self-medication against fatigue). A meta-analysis of clinical trials that looked at the effectiveness of nicotine patches as an aid to quit smoking, found that patch-users experienced more sleep disturbances than controls. The effects were proportional to the strength of the patch, and worse when it was left on overnight^18^, suggesting that nicotine has a causal, negative effect on sleep. In the other direction, smokers who were offered cigarettes or money picked cigarettes more often when they were sleep deprived, even when the monetary value was greater than that of the cigarette^19^. Finally, animal work has suggested that passive smoking, via an effect on gene expression, can alter circadian rhythm (which determines chronotype)^20^. Overall, current findings are mixed and have focussed on short-term rather than long-term effects. Novel methods are needed to fully disentangle the complex relationship between smoking and sleep. To distinguish genetic correlation from causal relationships, Mendelian randomization (MR) analysis can be applied. MR infers causality by taking a set of genetic variants robustly associated with an exposure variable as a proxy for this exposure and estimating its causal effect on an outcome variable^21^. Potential horizontal pleiotropy (genetic variants affecting the outcome directly, not acting through the exposure) can be assessed with sensitivity analyses.

In the present study, we first calculated genetic correlations between smoking (smoking initiation, cigarettes smoked per day and smoking cessation), and sleep behaviours (sleep duration, chronotype and insomnia) based on summery-level data of large, published GWA studies. We then employed MR – using individual level data of 333,581 participants of UK Biobank and summary-level data – to test bidirectional, causal effects between smoking and sleep behaviours.

## Methods

### Data sources

For LD score regression and MR with summary-level data (‘two-sample MR’) we used summary statistics from GWA studies on smoking initiation (*n*=74,035), cigarettes smoked per day (*n*=38,181) and smoking cessation (being a former versus a current smoker) (*n*=41,278)^14^, sleep duration (measured in hours and as undersleeping (≤6 hours versus 7-8 hours) and oversleeping (≥9 hours versus 7-8 hours)) and chronotype (measured on a five-point scale with 2 coded as being a ‘morning person’ and -2 being an ‘evening person’) (*n*=128,266)^15^ and insomnia (usually having trouble falling asleep at night or waking up in the middle of the night (‘cases’) versus never/rarely or sometimes having these problems (‘controls’)) (*n*=113,006)^16^.

For MR with individual level data (‘one-sample MR’), we obtained data of 333,581 participants of UK Biobank^22^. Details on quality control of the genetic data within the MRC-IEU are available in the Supplementary material.

## Statistical analysis

### LD Score Regression

Genetic correlations between smoking and sleep behaviours were estimated with LD-score regression^17^. This method is based on the expected relationship between the degree of linkage disequilibrium (LD) between single nucleotide polymorphisms (SNPs), and the strength of their association with the phenotype in question as derived from GWAS. SNPs that are in high LD with many neighbouring SNPs have a higher chance of tagging a causal genetic variant. From this it follows that SNPs with a higher degree of LD with neighbouring show larger effect sizes. This information is then used to compute a genetic correlation between two phenotypes. For a more detailed description of LD-score regression we refer to the work of Bulik-Sullivan and colleagues^17^.

### MR with individual level data

We investigated causal effects of *cigarettes smoked per day* and *smoking cessation* on sleep behaviours using MR with individual level data. Because genetic variants previously associated with cigarettes smoked per day and smoking cessation were identified in samples of smokers^14^, MR analyses investigating causal effects of these variables as exposures have to be stratified by smoking status. For cigarettes smoked per day, we used the genetic variant rs16969968 as instrumental variable – the minor allele of this SNP is associated with smoking, on average, one additional cigarette per day^23^. Analyses were performed in never, former and current smokers separately. For smoking cessation, we used the genetic variant rs3025343 as instrumental variable. The major allele of this variant has been associated with a higher odds of being a former versus a current smoker^14^.

Analyses were performed in ever smokers (current + former smokers) and never smokers separately. Associations between the genetic instruments and the exposure variables as well as common confounding variables are provided in the supplementary material (Supplementary Tables 1 to 7).

MR entailed linear and logistic regression analyses, performed in STATA. The genetic instrument for the smoking (exposure) variable was the independent variable (coded as 0, 1 or 2 risk alleles) while the sleep (outcome) variable was the dependent variable. If, for instance, the genetic instrument for smoking more cigarettes per day predicts sleep problems in smokers, but not in non-smokers, this would suggest a causal effect of smoking on sleep.

### Two sample MR with summary-level data

Next, we investigated causal effects for all other relationships between smoking and sleep, using MR with summary-level data^24^. In this approach, the gene-exposure association and the gene-outcome association are obtained from two different samples. Genetic instruments for the exposure variables were selected in the exposure GWAS at two levels of significance, first using SNPs reaching genome-wide significance (*p*<5 × 10^−8^) and second using a more liberal threshold of *p*<1 × 10^−5^ (independent SNPs were identified by pruning on *r^2^*<0.001). Gene-outcome associations, for SNPs associated with the exposure, were then extracted from the outcome GWAS. When SNPs were not available in the outcome GWAS, proxies were used (LD R^2^>0.8; identified using online tool SNiPa). A full list of the SNPs used in each analysis is provided in Supplementary Table 1.

Analyses were conducted using the R package of MR-Base, a database and analytical platform to perform MR^25^. Causal estimates were calculated using the Wald Ratio in case of genetic instruments consisting of a single SNP^24^. Where multiple SNPs were available, these were combined using Inverse-Variance Weighted fixed effects-meta-analysis (IVW)^26^. The IVW estimate is the mean average of the Wald ratios of all SNPs, inversely weighted by their standard error. Conveniently, two-sample MR allows sensitivity analyses that are more robust to horizontal pleiotropy, albeit less powerful. For genetic instruments that contained a sufficient number of SNPs (≥10), we employed two commonly used sensitivity analyses; weighted median regression and MR-Egger regression. Weighted median regression is able to provide a consistent estimate of a causal effect even when up to 50% of the weight in a polygenic score comes from invalid instruments^27^. MR-Egger is a variation of a test used in meta-analyses to test for small study bias. It rests on the InSIDE assumption (Instrument Strength Independent of Direct Effect) which means that the strength of the instrument should not correlate with the direct effect that the instrument has on the outcome and which is a much weaker assumption than that of no horizontal pleiotropy^28^.

## Results

### LD Score regression

The results of LD score regression are presented as a forest plot in **Figure 1**. We found strong evidence for positive genetic correlations between insomnia and cigarettes per day (*rg*=0.27, 95% CI 0.06 to 0.49) and insomnia and smoking initiation (*rg*=0.15, 95% CI 0.009 to 0.28), as was reported previously^16^. We also observed a positive genetic correlation between undersleeping and smoking initiation (*rg*=0.26, 95% CI 0.13 to 0.39) and, in agreement with this, a negative correlation between sleep duration and smoking initiation (*rg*=-0.14, 95% CI -0.26 to -0.01). The strongest correlation we observed was between undersleeping and cigarettes per day, *rg*=0.42 (95% CI 0.19 to 0.66), which, together with the other correlations described here, points to less or poorer sleep being associated with smoking (more heavily). Finally, we observed a negative correlation between chronotype and smoking cessation, *rg*=-0.18 (95% CI -0.31 to -0.06), indicating that genetic variants associated with being a morning person are also associated with reduced likelihood of quitting smoking. We found no clear evidence for genetic correlations between the other sleep and smoking behaviours.

**Figure 1.**
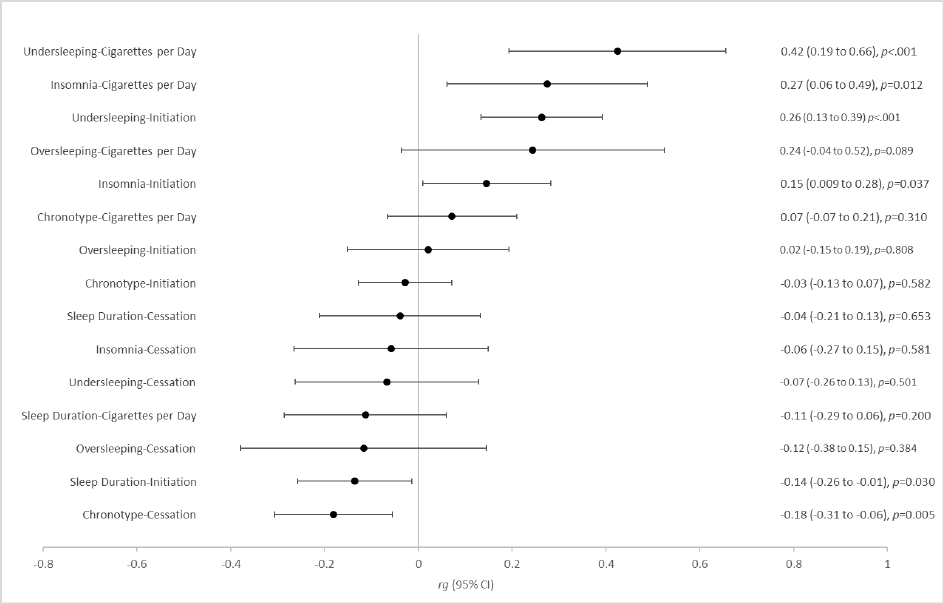
Forest plot depicting genetic correlations (*r*g), as calculated with LD score regression, between smoking behaviours and sleep behaviours

### MR with individual level data

The minor allele of rs16969968, which is associated with smoking more cigarettes per day was associated with a lower chronotype score (i.e., being more of an ‘evening’ person) in current smokers (beta=-0.062, 95% CI -0.084 to -0.039) (**Table 1**). A weaker, but similar effect was found in former smokers (beta=-0.016, 95% CI -0.027 to -0.005). In never smokers, the effect was in the opposite direction, with the smoking increasing allele of rs16969968 being associated with being a ‘morning’ person (beta=0.011, CI 0.003 to 0.020). There was no clear evidence for causal effects of smoking heaviness on sleep duration (in hours of sleep or measured as undersleeping or oversleeping) or insomnia, nor was there clear evidence for causal effects of smoking cessation on sleep behaviours.

**Table 1.** Mendelian randomisation (MR) analyses with individual level data, estimating causal effects of smoking behaviours (cigarettes per day and smoking cessation) on sleep behaviours (sleep duration, chronotype and insomnia).

| Exposure | Outcome | Smoking Status | N | beta/OR (95% CI) | p-value |
| --- | --- | --- | --- | --- | --- |
| rs16969968<br>(instrument for cigarettes per day) | Sleep duration | Never | 183,333 | -0.006 (-0.013 to 0.002) | 0.13 |
|  |  | Former | 117,743 | 0.009 (-0.001 to 0.018) | 0.07 |
|  |  | Current | 33,101 | -0.014 (-0.034 to 0.006) | 0.17 |
|  | Undersleeping | Never | 170,271 | 1.01 (0.99 to 1.03) | 0.12 |
|  |  | Former | 108,096 | 0.99 (0.97 to 1.01) | 0.24 |
|  |  | Current | 30,237 | 1.02 (0.98 to 1.05) | 0.37 |
|  | Oversleeping | Never | 141,833 | 0.99 (0.96 to 1.01) | 0.30 |
|  |  | Former | 89,716 | 1.01 (0.98 to 1.05) | 0.41 |
|  |  | Current | 23,486 | 0.98 (0.93 to 1.04) | 0.58 |
|  | Chronotype | Never | 183,925 | 0.011 (0.003 to 0.020) | 0.01 |
|  |  | Former | 118,041 | -0.016 (-0.027 to -0.005) | 0.004 |
|  |  | Current | 33,285 | -0.062 (-0.084 to -0.039) | <0.001 |
|  | Insomnia | Never | 184,184 | 0.99 (0.98 to 1.01) | 0.38 |
|  |  | Former | 118,181 | 1.01 (0.99 to 1.02) | 0.56 |
|  |  | Current | 33,343 | 1.00 (0.97 to 1.04) | 0.79 |
| rs3025343<br>(instrument for smoking cessation) | Sleep duration | Never | 183,333 | -0.007 (-0.018 to 0.003) | 0.18 |
|  |  | Ever | 117,743 | 0.005 (-0.009 to 0.018) | 0.52 |
|  | Undersleeping | Never | 170,271 | 1.01 (0.99 to 1.04) | 0.34 |
|  |  | Ever | 108,096 | 1.01 (0.98 to 1.04) | 0.47 |
|  | Oversleeping | Never | 141,833 | 0.99 (0.95 to 1.03) | 0.54 |
|  |  | Ever | 89,716 | 1.04 (0.99 to 1.09) | 0.11 |
|  | Chronotype | Never | 183,925 | -0.002 (-0.015 to 0.010) | 0.72 |
|  |  | Ever | 118,041 | 0.011 (-0.006 to 0.027) | 0.20 |
|  | Insomnia | Never | 183,082 | 1.01 (0.99 to 1.03) | 0.37 |
|  |  | Ever | 118,181 | 1.00 (0.98 to 1.03) | 0.85 |

For cigarettes per day coefficients represent the change in outcome per additional minor allele of rs16969968. For smoking cessation coefficients represent the change in outcome per additional major allele of rs3025343. Sleep duration was measured in hours per day. Undersleeping was defined as sleeping ≤6 hours versus 7-8 hours and oversleeping as ≥9 hours versus 7-8 hours. Chronotype was measured on a five-point scale, with higher scores indicating being more of a 'morning person' and lower scores more of an 'evening person'. Insomnia was measured as usually having trouble falling asleep at night or waking up in the middle of the night ('cases') versus never/rarely or sometimes having these problems ('controls').

### Two sample MR with summary-level data

Using MR with summary-level data of published GWAS studies on smoking and sleep behaviours, there was no clear evidence for a causal influence of smoking initiation on sleep duration, chronotype or insomnia (**Table 2**).

**Table 2.**
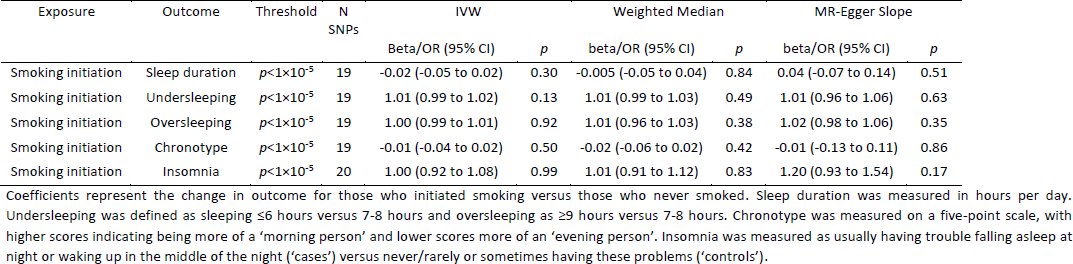
Two-sample Mendelian randomisation (MR) analyses with summary-level data, estimating causal effects of smoking initiation on sleep behaviours (sleep duration, chronotype and insomnia).

In the other direction, there was no clear evidence for causal effects of sleep duration or chronotype on smoking behaviour (**Table 3**). There was some weak evidence for insomnia increasing the number of cigarettes smoked per day, but only with the genetic instrument combining 16 SNPs (IVW beta=1.21, 95% CI=0.20 to 2.22). MR-Egger regression showed an opposite direction of effect (IVW beta=-1.07, 95% CI=-5.32 to 3.18), but with wide confidence intervals. There was also weak evidence that insomnia causally influences smoking cessation such that having insomnia complaints decreases the odds of being a former versus a current smoker (IVW OR 0.80 95% CI=0.65 to 0.97). Similar directions of effect were found with both weighted median (IVW OR=0.86 95% CI=0.66 to 1.13) and MR-Egger regression (IVW OR=0.49 95% CI=0.21 to 1.11) methods. There was no clear evidence for directional pleiotropy from insomnia to smoking cessation, as indicated by the Egger intercept (IVW OR=1.03, 95% CI=0.98, 1.07; Supplementary Table 5), nor was there evidence for heterogeneity (Cochran’s Q for IVW=14.0, *p*=0.526); Supplementary Table 4). For all other relationships tested, there was no clear evidence for causality.

**Table 3.**
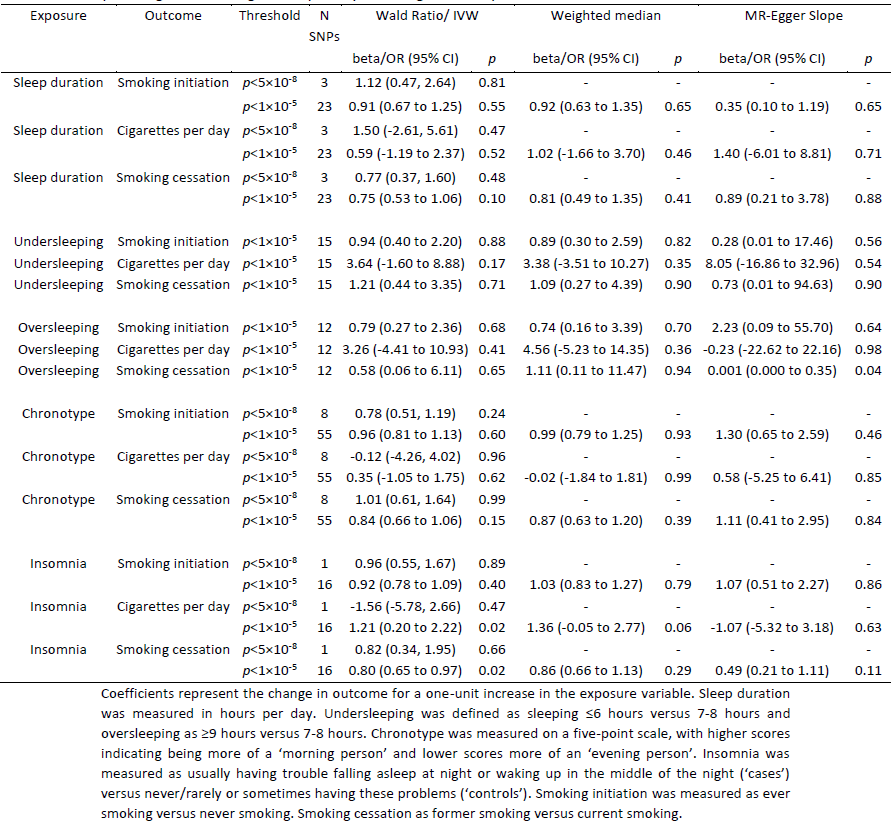
Two-sample Mendelian randomisation (MR) analyses with summary-level data, estimating causal effects of sleep behaviours (sleep duration, chronotype and insomnia) on smoking behaviours (smoking initiation, cigarettes per day, smoking cessation).

## Discussion

We found evidence of moderate genetic correlations between smoking and sleep behaviours. Genetic risk variants that were associated with insomnia also increased the odds of initiating smoking and were associated with smoking more cigarettes per day. Consistent with this finding, genetic variants that predicted shorter sleep duration increased the odds of initiating smoking. Finally, genetic variants that made it more likely to be a morning person (chronotype) were associated with a lower odds of smoking cessation. Possible causal relationships underlying these genetic correlations were tested with Mendelian randomization (MR) analyses. Using MR with individual level data, we found compelling evidence that smoking more cigarettes per day decreases the odds of being a morning person. MR with summary-level data indicated weak evidence for insomnia causally increasing the number of cigarettes smoked and decreasing the odds of quitting smoking.

While a genetic variant that is robustly associated with smoking heaviness was associated with a *lower* likelihood of being a morning person in smokers – consistent with a causal effect of smoking heaviness on chronotype – this same genetic variant was associated with a *higher* likelihood of morningness in never smokers. This suggests that there is pleiotropy, such that the genetic instrument plays a (direct) role in chronotype. However, given that the direction of effect in never smokers is opposite to that in current smokers, and in former smokers the effect is in the same direction as smokers but less strong, it is unlikely that pleiotropy is driving the association we see in smokers. This suggests that previously reported observational associations between smoking, nicotine dependence and being an evening person^7^ may at least in part be explained by a causal effect of smoking on chronotype. One explanation for this is that the psychoactive properties of nicotine allow individuals to stay more alert into the evening^7^. There is also evidence from animal research that both nicotine and nicotinic acetylcholine receptors (nAChRs) play a role in the circadian system^20,29^. This could explain our finding of an association in never smokers, given that the genetic variant, rs16969968, is a missense mutation which causes a functional change in the α5 nAChR subunit protein^30^.

MR with summary-level data provided suggestive evidence that insomnia causally increases smoking heaviness. While the evidence was weak, it was supported by a considerable genetic correlation between insomnia and cigarettes per day (*r*g=0.27). In line with this finding, there was weak evidence that insomnia decreases the odds of smoking cessation. One mechanism underlying a causal relationship from insomnia to smoking heaviness and cessation could be that smokers self-medicate against fatigue with cigarettes^19^. If smoking cigarettes alleviates the negative consequences of insomnia, this could make it harder for smokers with insomnia to give up smoking. We didn’t find similar evidence for causal effects of sleep duration (in hours or as undersleeping) on smoking behaviour, which may seem unexpected. However, insomnia is not necessarily characterized by a short *total* sleep duration^31^ and individuals with a naturally short duration of sleep may not experience fatigue. Our suggestive findings that insomnia causally increases smoking heaviness and decrease the odds of quitting can be informative for developing strategies to improve smoking cessation success. Treatments that are currently offered (nicotine replacement therapies and medications such as bupropion or varenicline) are only moderately effective^32^ and insomnia may be a promising novel target that could complement existing treatments. This was also highlighted in a recent review paper, which reported extensive conceptual support for sleep therapy as an adjunctive treatment for smoking^33^.

Given the large sample sizes employed for our analyses, a major strength to our study is that we had much power to detect small effects. By combining multiple methods – LD score regression, MR with individual level data and with summary-level data – we were able to differentiate shared genetic risk factors between smoking and sleep, from possible casual effects. A limitation to our study is that for some of the phenotypes (smoking initiation, undersleeping, oversleeping) we could only test effects with a genetic instrument that included SNPs under the threshold *p*<1×10^−5^. These instruments are less robustly associated with the exposure variable and therefore less precise when testing causal effects on an outcome variable.

In summary, our findings suggest that smoking and sleep behaviours are genetically correlated and that some of this correlation reflects causal effects. Smoking more cigarettes per day seems to affect circadian rhythm (shifting it to being more of an ‘evening person’) while in the other direction we found weak evidence that insomnia increases smoking heaviness and decreases the odds of successfully quitting smoking. These findings increase our knowledge of the complex relationship between smoking and sleep and can help develop more evidence-based treatments for nicotine dependence by focussing on insomnia as a novel treatment target.

## Acknowledgements

None.

## References

1. Wetter DW, Young TB. The relation between cigarette smoking and sleep disturbance. Prev Med. 1994;23(3):328–334. doi:10.1006/pmed.1994.1046.

2. Åkerstedt T, Narusyte J, Alexanderson K, Svedberg P. Sleep Duration, Mortality, and Heredity—A Prospective Twin Study. Sleep. 2017;40(10):zsx135–zsx135. doi:10.1093/sleep/zsx135.

3. Boakye D, Wyse CA, Morales-Celis CA, et al. Tobacco exposure and sleep disturbance in 498 208 UK Biobank participants. J Public Health (Bangkok). 2017:1–10. doi:10.1093/pubmed/fdx102.

4. Nguyen-Louie TT, Brumback T, Worley MJ, et al. Effects of sleep on substance use in adolescents: a longitudinal perspective. Addict Biol. 2017. doi:10.1111/adb.12519.

5. Patterson F, Grandner MA, Lozano A, Satti A, Ma G. Transitioning from adequate to inadequate sleep duration associated with higher smoking rate and greater nicotine dependence in a population sample. Addict Behav. 2018;77(Supplement C):47–50. doi:https://doi.org/10.1016/j.addbeh.2017.09.011.

6. Peltier MR, Lee J, Ma P, Businelle MS, Kendzor DE. The influence of sleep quality on smoking cessation in socioeconomically disadvantaged adults. Addict Behav. 2017;66:7–12. doi:10.1016/j.addbeh.2016.11.004.

7. Broms U, Pennanen M, Patja K, et al. Diurnal Evening Type is Associated with Current Smoking, Nicotine Dependence and Nicotine Intake in the Population Based National FINRISK 2007 Study. J Addict Res Ther. 2012;S2. https://www.ncbi.nlm.nih.gov/pubmed/22905332.

8. Khan MS, Aouad R. The Effects of Insomnia and Sleep Loss on Cardiovascular Disease. Sleep Med Clin. 2017;12(2):167–177. doi:https://doi.org/10.1016/j.jsmc.2017.01.005.

9. Services USD of H and H. The Health Consequences of Smoking - 50 Years of Progress: A Report of the Surgeon General. Atlanta: U.S. Department of Health and Human Services, Centers for Disease Control and Prevention, National Center for Chronic Disease Prevention and Health Promotion, Office on Smoking and Health, 2014; 2014.

10. Businelle MS, Kendzor DE, Reitzel LR, et al. Pathways linking socioeconomic status and postpartum smoking relapse. Ann Behav Med. 2013;45(2):180–191. doi:10.1007/s12160-012-9434-x.

11. Grandner MA, Patel NP, Gehrman PR, et al. Who gets the best sleep? Ethnic and socioeconomic factors related to sleep complaints. Sleep Med. 2010;11(5):470–478. doi:10.1016/j.sleep.2009.10.006.

12. Lind MJ, Aggen SH, Kirkpatrick RM, Kendler KS, Amstadter AB. A Longitudinal Twin Study of Insomnia Symptoms in Adults. Sleep. 2015;38(9):1423–1430. doi:10.5665/sleep.4982.

13. Vink JM, Willemsen G, Boomsma DI. Heritability of Smoking Initiation and Nicotine Dependence. Behav Genet. 2005;35(4):397–406. doi:10.1007/s10519-004-1327-8.

14. Furberg H, Kim Y, Dackor J, et al. Genome-wide meta-analyses identify multiple loci associated with smoking behavior. Nat Genet. 2010;42(5):441–447. doi:10.1038/ng.571.

15. Jones SE, Tyrrell J, Wood AR, et al. Genome-Wide Association Analyses in 128,266 Individuals Identifies New Morningness and Sleep Duration Loci. PLoS Genet. 2016;12(8):e1006125. doi:10.1371/journal.pgen.1006125.

16. Hammerschlag AR, Stringer S, de Leeuw CA, et al. Genome-wide association analysis of insomnia complaints identifies risk genes and genetic overlap with psychiatric and metabolic traits. Nat Genet. 2017. doi:10.1038/ng.3888.

17. Bulik-Sullivan B, Finucane HK, Anttila V, et al. An atlas of genetic correlations across human diseases and traits. Nat Genet. 2015;47(11):1236–1241. doi:10.1038/ng.3406.

18. Greenland S, Satterfield MH, Lanes SF. A meta-analysis to assess the incidence of adverse effects associated with the transdermal nicotine patch. Drug Saf. 1998;18(4):297–308. https://www.ncbi.nlm.nih.gov/pubmed/9565740.

19. Hamidovic A, de Wit H. Sleep deprivation increases cigarette smoking. Pharmacol Biochem Behav. 2009;93(3):263–269. doi:10.1016/j.pbb.2008.12.005.

20. Numaguchi S, Esumi M, Sakamoto M, et al. Passive cigarette smoking changes the circadian rhythm of clock genes in rat intervertebral discs. J Orthop Res. 2016;34(1):39–47. doi:10.1002/jor.22941.

21. Davey Smith G, Ebrahim S. “Mendelian randomization”: can genetic epidemiology contribute to understanding environmental determinants of disease?*. Int J Epidemiol. 2003;32(1):1–22. doi:10.1093/ije/dyg070.

22. Thompson SG, Willeit P. UK Biobank comes of age. Lancet. 2015;386(9993):509–510. doi:10.1016/S0140-6736(15)60578-5.

23. Ware JJ, van den Bree MB, Munafo MR. Association of the CHRNA5-A3-B4 gene cluster with heaviness of smoking: a meta-analysis. Nicotine Tob Res. 2011;13(12):1167–1175. doi:10.1093/ntr/ntr118.

24. Burgess S, Scott RA, Timpson NJ, Davey Smith G, Thompson SG, EPIC- InterAct Consortium. Using published data in Mendelian randomization: a blueprint for efficient identification of causal risk factors. Eur J Epidemiol. 2015;30(7):543–552. doi:10.1007/s10654-015-0011-z.

25. Zheng J, Haycock P, Hemani G, et al. LD hub and MR-base: online platforms for preforming LD score regression and Mendelian randomization analysis using GWAS summary data. Behav Genet. 2016;46(6):815.

26. Lawlor DA, Harbord RM, Sterne JAC, Timpson N, Davey Smith G. Mendelian randomization: Using genes as instruments for making causal inferences in epidemiology. Stat Med. 2008;27(8):1133–1163. doi:10.1002/sim.3034.

27. Bowden J, Davey Smith G, Haycock PC, Burgess S. Consistent Estimation in Mendelian Randomization with Some Invalid Instruments Using a Weighted Median Estimator. Genet Epidemiol. 2016;40(4):304–314. doi:10.1002/gepi.21965.

28. Bowden J, Davey Smith G, Burgess S. Mendelian randomization with invalid instruments: effect estimation and bias detection through Egger regression. Int J Epidemiol. 2015;44(2):512–525. doi:10.1093/ije/dyv080.

29. O’Hara BF, Edgar DM, Cao VH, et al. Nicotine and nicotinic receptors in the circadian system. Psychoneuroendocrinology. 1998;23(2):161–173. doi:https://doi.org/10.1016/S0306-4530(97)00077-2.

30. Lassi G, Taylor AE, Timpson NJ, et al. The CHRNA5-A3-B4 Gene Cluster and Smoking: From Discovery to Therapeutics. Trends Neurosci. 2016;39(12):851–861. doi:10.1016/j.tins.2016.10.005.

31. Johann AF, Hertenstein E, Kyle SD, et al. Insomnia with objective short sleep duration is associated with longer duration of insomnia in the Freiburg Insomnia Cohort compared to insomnia with normal sleep duration, but not with hypertension. Romigi A, ed. PLoS One. 2017;12(7):e0180339. doi:10.1371/journal.pone.0180339.

32. Ferguson J, Bauld L, Chesterman J, Judge K. The English smoking treatment services: one-year outcomes. Addiction. 2005;100(s2):59–69. doi:10.1111/j.1360-0443.2005.01028.x.

33. Patterson F, Grandner MA, Malone SK, Rizzo A, Davey A, Edwards DG. Sleep as a Target for Optimized Response to Smoking Cessation Treatment. Nicotine Tob Res. October 2017. doi:10.1093/ntr/ntx236.

